# Base-edited CAR T Cells for combinational therapy against T cell malignancies

**DOI:** 10.1101/2020.07.30.228429

**Authors:** Christos Georgiadis, Jane Rasaiyaah, Soragia Athina Gkazi, Roland Preece, Aniekan Etuk, Abraham Christi, Waseem Qasim

## Abstract

Targeting T cell malignancies using chimeric antigen receptor (CAR) T cells is hindered by ‘T v T’ fratricide against shared antigens such as CD3 and CD7. Genome-editing can overcome such hurdles through targeted disruption of problematic shared antigens. Base editing offers the possibility of seamless disruption of gene expression through the creation of stop codons or elimination of splice donor or acceptor sites. We describe the generation of fratricide resistant, T cells by orderly removal of shared antigens such as TCR/CD3 and CD7 ahead of lentiviral mediated expression of CARs specific for CD3 or CD7. Molecular interrogation of base edited cells confirmed virtual elimination of chromosomal translocation events detected in conventional Cas9 treated cells. Interestingly, co-culture of 3CAR and 7CAR cells resulted in ‘self-enrichment’ yielding populations that were 99.6% TCR-/CD3/^-^CD7^-^. 3CAR or 7CAR cells were able to exert specific cytotoxicity against their relevant target antigen in leukaemia lines with defined CD3 and/or CD7 expression as well as primary T-ALL cells. Co-cultured 3CAR/7CAR cells exhibited the highest level of cytotoxicity against T-ALL targets expressing both target *in vitro* and an *in vivo* human:murine chimeric model. While APOBEC editors can reportedly exhibit guide-independent deamination of both DNA and RNA, we found no evidence of promiscuous base conversion activity affecting CAR antigen specific binding regions which may otherwise redirect T cell specificity. Combinational infusion of fratricide resistant anti-T CAR T cells may enable enhanced molecular remission ahead of allogeneic haematopoietic stem cell transplantation for T cell malignancies.

## Introduction

T cell acute lymphoblastic leukaemia (T-ALL) arises from lymphoid precursors and is typically associated with poorer prognosis to comparable B cell leukaemia. Standard therapy for T-ALL, uses high intensity multi-agent chemotherapy where tolerated and sensitive minimal residual disease (MRD) assessments can determine when molecular remission is achieved. However, 5 year survival for T-ALL is < 75% for children and <50% for adults (1) and while salvage therapy such as nelarabine (2) and allogeneic stem cell transplantation (SCT) (3) are widely used, innovative therapies have lagged behind progress for other leukaemias. Strategies using chimeric antigen receptor (CAR) T cells are being widely applied against B cell malignancies, and targeting of surface antigens such as CD19, CD20 and CD22, which are also present on healthy B cells, has been shown to be broadly tolerable (4-6). A proportion of patients with sustained B cell aplasia require long term immunoglobulin replacement, and some may go on to allogeneic transplant and donor derived immunological recovery (7, 8). Similar approaches to tackle T cell malignancies have been more challenging because surface expression of surface antigens such as the TCRαβ/CD3 complex results in compromising fratricidal effects during T cell production. We have previously reported how this issue can be circumvented by removal of cell surface CD3 through disruption of TRAC expression by TALEN genome editing (9). Therapeutic applications envisaged strictly time-limited effector activity to secure remission, and then rapid reconstitution of a diverse T cell compartment given that substitution of T cell function (unlike B cell function) is not possible. The kinetics of autologous T cell recovery in such settings has yet to be defined but could take many months and may be limited to post thymic populations with restricted TCR repertoire diversity. The availability of a Human Leukocyte Antigen (HLA)-matched allogeneic donor may offer the prospect of donor derived T cell recovery, but in either setting the strategy relies on first securing rapid and deep molecular remission through CAR T effects. The additional issue of antigen escape or incomplete expression on disease populations has resulted in a quest for additional antigens, and proposals to target multiple antigens simultaneously. Similar approaches are under development for B cell malignancies, for example by using CAR T cells targeting CD19 and CD22 or CD123 generated using bicistronic constructs or tandem CAR configurations (10-12). Alternatively, a more physiological and flexible strategy involves the generation of multiple T cell banks with differing CAR specificities to be used in combination, as determined by particular disease specific cell surface molecule expression. We have investigated CAR T cells specific for CD3 and CD7 for combinational use against T cell malignancies. The latter is a 40-kD single-domain Ig superfamily transmembrane glycoprotein encoded on chromosome 17q25.3 and highly expressed on normal and malignant T cells. Others have previously generated anti-CD7 CAR T cells following expression of inhibitory proteins (13) or CRISPR/Cas9 (14, 15) editing of T cells to avoid fratricidal effects. We addressed the risk of fratricide and cross-recognition during co-effector activity by removal of cell surface targets recognised by the respective CARs. To achieve this efficiently, and at reduced risk of genotoxicity, multiplexed base editing was combined with lentiviral CAR delivery. Base editing used CRISPR guided chemical deamination for highly precise, seamless, cytidine to uridine to thymidine (C→U→T) conversion which then directed the creation of stop codons or disrupted splice donor/acceptor sites. The technology used BE3 (16), comprising a deactivated Cas9 nickase fused to rat APOBEC1 (the deaminase) and a single uracyl glycosylase inhibitor (UGI) delivered as mRNA by electroporation in combination with CRISPR guides. Highly specific RNA guided cytidine deamination in T cells ahead of lentiviral transduction ensured target antigen disruption before CAR expression. An important consideration in the context of multiplexed editing to date has been the creation of chromosomal translocations following nuclease-mediated double strand breakage, especially when multiple genomic loci are targeted simultaneously. Previously, we had reported that around 5% of TALEN edited CAR19 T cells, disrupted at both TRAC and CD52 loci, exhibited karyotypic abnormalities (17, 18), and a similar frequency was recently reported in CRISPR/Cas9 edited T cells expressing recombinant TCR (19). Advantages of base editing were expected to include high efficiency editing (20) and a notable reduction of otherwise predictable translocations between edited chromosomes. We report that removal of shared antigens permits generation of anti-CD3 and anti-CD7 CAR T cells and enables combinational effector activity against T cell targets. The strategies are readily scalable through the adaption of existing semi-automated manufacturing processes. It is envisaged that a time limited therapeutic application of combinational anti-T CAR T cells could deliver deep molecular remission ahead of conditioning and programmed allogeneic SCT in difficult to treat T-ALL.

## Results

### Combinational CAR T cell targeting of CD3 and CD7 requires multiplexed base conversion for simultaneous removal of shared antigens

We have previously described a chimeric antigen receptor, 3CAR, specific for CD3ε comprising a scFv sequence derived from the therapeutic monoclonal antibody OKT3 fused to a CD8 stalk and 41BB-CD3ζ second generation CAR (9). TALEN mediated genetic disruption of the TRAC locus ensured that T cells subsequently transduced with a lentiviral-CD3 CAR targeting vector lacked cell surface expression of TCRαβ/CD3 and thereby evaded fratricidal effects. Similar effects were readily elicited using CRISPR directed Cas9 editing, and investigations revealed the critical importance of orderly disruption of the target antigen locus ahead of CAR delivery and expression. The same principles were applied to evaluate a CAR specific for CD7 **(Figure 1A-C)**. The construct was generated using light and heavy variable domains derived from the 3A1e hybridoma as previously reported (15) and fused to a CD8TM-41BB-CD3ζ configuration within a third generation SIN lentiviral vector **(Figure 1B)**. Of note, in initial experiments, we uncovered CD7 epitope masking effects when T cells were transduced to express this particular CAR, probably as a result of the scFv binding CD7 on the cell surface. Thus, CD7 expression in non-edited T cells was only detectable by staining with an anti-CD7 monoclonal antibody distinct from 3A1e because of competitive masking by CAR7 **(Supplementary Figure 1)**.

**Figure 1.**
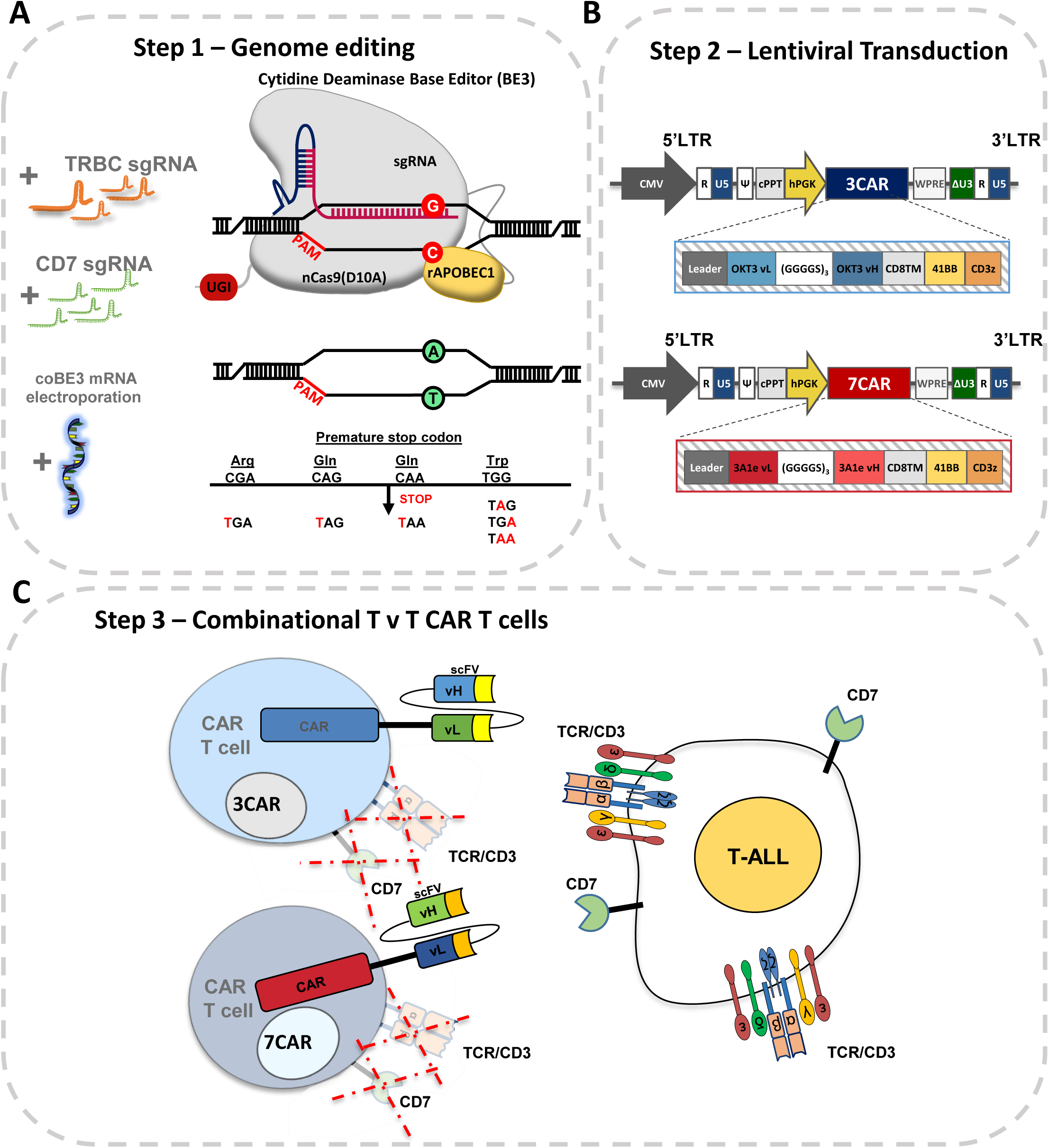
Generation of ‘T v T’ fratricide resistant CAR T cells. **A)** Schema of base editing for T cells employing 3^rd^ generation codon optimised cytidine base deaminase (coBE3) fused to deactivated D10A Cas9 nickase and uracil glycosylase inhibitor (UGI) delivered as mRNA along with TRBC and CD7 single guide RNA (sgRNA). C->U->T conversion (G->A antisense strand) resulting in STOP codon. **B)** Lentiviral transduction of edited cells from step 1 using 3^rd^ generation lentiviral vectors. Lentiviral plasmid configuration of CD3ε targeting 2^nd^ generation chimeric antigen receptor comprising OKT3 vL and vH scFv sequence fused to CD8 transmembrane domain (TM), 41BB co-stimulatory and CD3z activation domains under the control of a hPGK promoter. Lentiviral plasmid configuration of CD7 targeting 2^nd^ generation CAR comprising 3A1e vL and vH scFv sequence fused to CD8TM-41BB-CD3z under the control of a hPGK. **C)** coBE3 edited T cells devoid of shared antigens TCR/CD3 and CD7 surface receptors expressing either 3CAR or 7CAR evade fratricide and target T-ALL. BE: base editor; APOBEC: (apolipoprotein B mRNA editing enzyme, catalytic polypeptide-like); sgRNA: single guide RNA; PAM: protospacer adjacent motif; LTR: long terminal repeat; CMV: cytomegalovirus promoter; CAR: chimeric antigen receptor; cPPT: central polypurine track; U5: untranslated 5’ region; DU3: delta untranslated 3’ region; hPGK: human phosphoglycerate kinase promoter; vL: variable light chain; vH: variable heavy chain.

Genome editing at multiple loci was applied to prevent fratricide during co-culture using sgRNA guides against TRBC1/2 and CD7 capable of operating with both CRISPR/Cas9 and CBE reagents **(Figure 1C)**. Thus stabilised guides with 2’-O-methyl 3’ phosphorothioate modifications were electroporated along with CleanCap spCas9 mRNA approximately 24 hours after activation with anti-CD3/CD28 antibodies.

Transient RNA mediated expression of editing reagents was designed to ensure dilutional elimination in dividing cells for reduced off-target effects and immunogenicity. After a further 24 hours, lentiviral 3CAR or 7CAR transduction (MOI 5) was followed by 12 days of culture. High level surface disruption of TCRαβ/CD3 and CD7 by flow cytometry (63.7% and 60.8%, respectively) was achieved using SpCas9 mRNA and was associated with robust CAR expression throughout culture (78.2% 3CAR^+^ and 80.5% 7CAR^+^) **(Supplementary Figure 2A).** Application of coBE3 mRNA mediated similar levels of disruption of cell surface expression of TCRαβ/CD3 (63.4%) and CD7 (49.3%) with double knockout accounting for 37.5% of cells **(Figure 2A & B)**. This was verified at the molecular level as 61% and 47% G>A conversion at positions G_5_ G_6_ in TRBC and 61% conversion of C>T at positions C_8_ in CD7 **(Figure 2C, D & E)**. For TRBC, ≤6% G>A conversions occurred outside the expected BE3 editing window and ≤9% C>T for CD7 **(Figure 2 D & E)**. Neither SpCas9 editing nor cytidine deamination impacted on CAR expression with comparably high level 3CAR (SpCas9: 78.2%; coBE3: 53.1%) and 7CAR (SpCas9: 80.5%; coBE3: 70.1% expression at the end of production **(Figure 2A & B)**. Co-culture of edited 3CAR and 7CAR products demonstrated the critical advantage of shared antigen removal from both products yielding a 99.6% TCRαβ^-^CD7^-^ population expressing high levels of 3CAR/7CAR (SpCas9: 78.4%; coBE3: 80.9%) suggesting evasion of fratricide effects **(Figure 2A Supplementary Figure 2A)**. Interestingly, co-culture also allowed for self-enrichment effects by eliminating residual CD7^+^ or TCRαβ^+^ cells, although further TCRαβ magnetic column-based depletion may be warranted to ensure stringent depletion of residual TCRαβ cells in the allogeneic setting **(Figure 2A & B & Supplementary Figure 2A)**.

**Figure 2.**
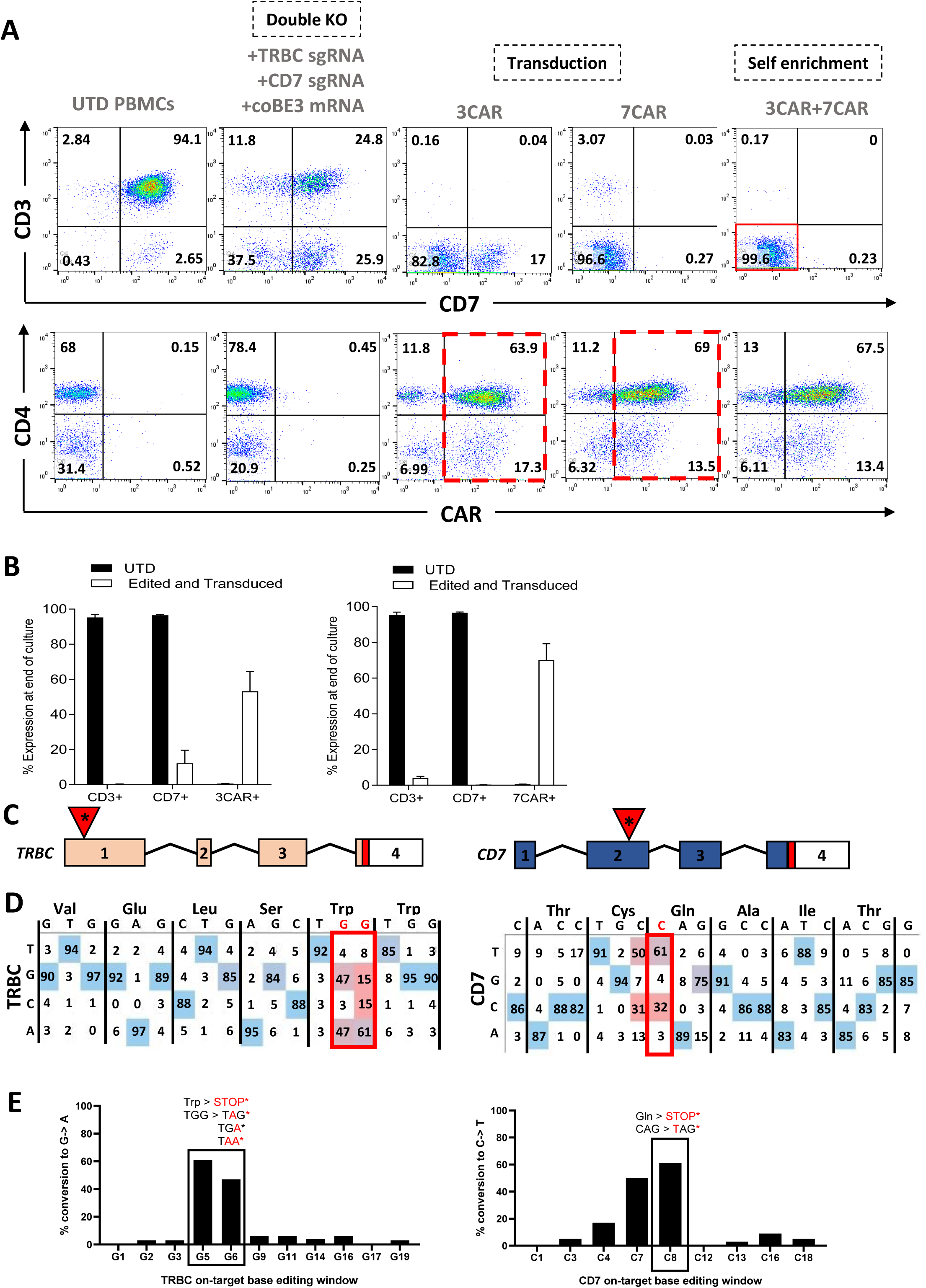
3CAR and 7CAR primary T cells evade fratricide during production. **A)** Phenotypic analysis of surface antigen CD3 and CD7 expression (top panel) and CAR expression (bottom panel) of CD3/CD28 activated peripheral blood mononuclear cells elcetroporated with sgRNA targeting TRBC and CD7 alongside coBE3 mRNA and subsequently transduced with 3CAR or 7CAR lentiviral vectors at MOI 5. Reduced TCR/CD3 and CD7 expression in edited groups and high level CAR expression with fratricide evasion (dotted red outline) was exhibited (n=4). Self-enrichment effects followed co-culture of 3CAR and 7CAR products (n=2) resulted in enriched TCR^-^CD7^-^ 3CAR/7CAR cells (red box). **B)** Proportion of CD3 or CD7 surface antigen and CAR expression at end of 3CAR (n=4), 7CAR (n=4) production or untransduced (UTD) (n=4) cells. Error bars represent SEM across (n=4) donors. **C)** Schematic of exonic regions within *TRBC* and *CD7* genes. Red marking in exons 4 of *TRBC* and *CD7* represent genomic translation stop sites followed by 5’ untranslated regions (white boxes). Red triangles with asterisk indicate position of base conversion resulting in premature stop codon formation. **D)** Representative base EDITR output of Sanger sequencing results from mixed 3CAR/7CAR co-culture DNA PCR amplicons of *TRBC* and *CD7* genomic loci. Sites of intended base conversion highlighted in red boxes. High frequency G->A (antisense) and C->T changes within editing window highlighted by red vs blue colour. **E)** Percentage of G->A conversions throughout TRBC-targeting protospacer sequence (left), and C->T conversions throughout CD7-targeting protospacer sequence (right).

### Cytidine base editing reduces predicted dsDNA chromosomal translocations

DSBs can lead to chromosomal translocations between sites of multiplexed editing, and this has been reported in TALEN and CRISPR/Cas9 modified T cells. Base conversion offers the prospect of seamlessly disrupting *TRBC1/2* and *CD7* without DSBs and thus we examined genomic DNA at the end of production 3CAR and 7CAR primary T cells following multiplexed TRBC1/2 and CD7 editing and compared the frequency of predicted translocations generated by spCas9 or coBE3 activity. DNA fragments bearing each of four predicted fusions between two disrupted chromosomal loci were synthesised to provide control readings for PCR reactions across the novel junctions **(Figure 3A)**. Using a combination of TRBC1/2 and CD7 binding primers, PCR, was able to detect signals in 3/4 sites for spCas9 (T1, T2, T3) and 1/4 (T1) sites for coBE3 treated samples **(Figure 3B)**. Upon further investigation, ddPCR quantified translocations detected for spCas9 edited samples at 0.25%-0.98% across all four combinations **(Figure 3C)** whereas only very low level events were detected for T1 and T3 in coBE3 edited indicating frequencies of <0.18% **(Figure 3C)**.

**Figure 3.**
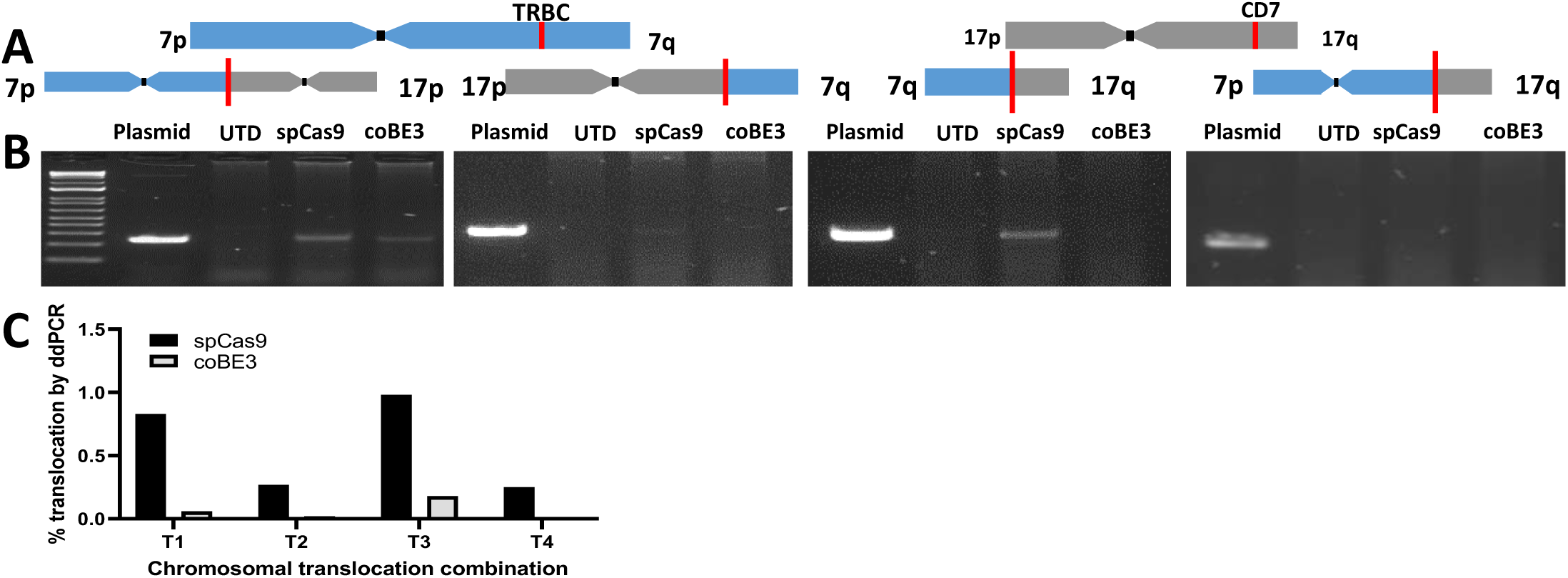
Multiplexed cytidine deamination reduces frequency of dsDNA break-mediated chromosomal translocations. **A)** Schematic of *TRBC* locus within chromosome 7 q-arm and *CD7* locus within chromosome 17 q-arm highlighted by red line. Four predicted chromosomal translocations generated following simultaneous dsDNA-mediated cleavage at *TRBC* and *CD7* loci. **B)** Gel electrophoresis of DNA products from either untransduced (UTD) or mixed 3CAR/7CAR co-cultures edited with spCas9 or coBE3 mRNA following PCR amplification with *TRBC* Fwd - *CD7* Fwd, *TRBC* Rev – *CD7* Fwd, *TRBC* Rev – *CD7* Rev and *TRBC* Fwd-*CD7* Rev primer combinations. Positive bands detected at ∼250bp. Control bands are PCR amplicons from of synthesised fusions. **C)** Histogram showing percentage of digital droplet PCR (ddPCR)-based quantification of four possible predicted translocations (T1-T4) in DNA from mixed 3CAR/7CAR co- cultures edited with spCas9 or coBE3 mRNA.

### 3CAR & 7CAR T cell cytotoxicity

*In vitro* function of 3CAR and 7CAR effector T cells was initially assessed by co-culture experiments with ^51^Cr labelled Jurkat T cell targets, either alone or in combination. Alone, 3CAR or 7CAR primary T cells exhibited high levels of cytotoxic activity against CD3^+^CD7^+^ targets, with both 3CAR and 7CAR effectors lysing >40% of targets at an E:T of 10:1 **(Figure 4A)**. Cytotoxic activity was comparable to spCas9-edited 3CAR or 7CAR cells indicating intact functional integrity after base editing **(Supplementary Figure 2C)**. Against antigen-negative controls, co-culture of either 3CAR or 7CAR effectors against CD3^-^CD7^-^ targets at an equivalent E:T ratio, resulted in <5% cytotoxicity. Specificity was further corroborated by co-culture of either 3CAR or 7CAR effectors with mixed population target cells, comprising CD3^+^CD7^-^ or CD3^-^CD7^+^ populations. This demonstrated CAR-specific clearance was restricted to cells exhibiting their respective antigens, and absence of lysis of antigen negative cell fractions **(Figure 4A)**. In a clinical setting, we envisage combined infusions of 3CAR and 7CAR T cells will offer the greatest prospect of leukaemic clearance. To model this, 3CAR T cells were co-cultured with 7CAR T cells and then subjected to potency testing in the same cytotoxicity assay **(Figure 4A)**. In combination, 3CAR/7CAR cells displayed highest cytotoxicity against antigen mixed CD3^-^CD7^+^ and CD3^+^CD7^-^ targets with ∼40% lysis at E:T ranges between 10:1 and 5:1 **(Figure 4A)**.

**Figure 4.**
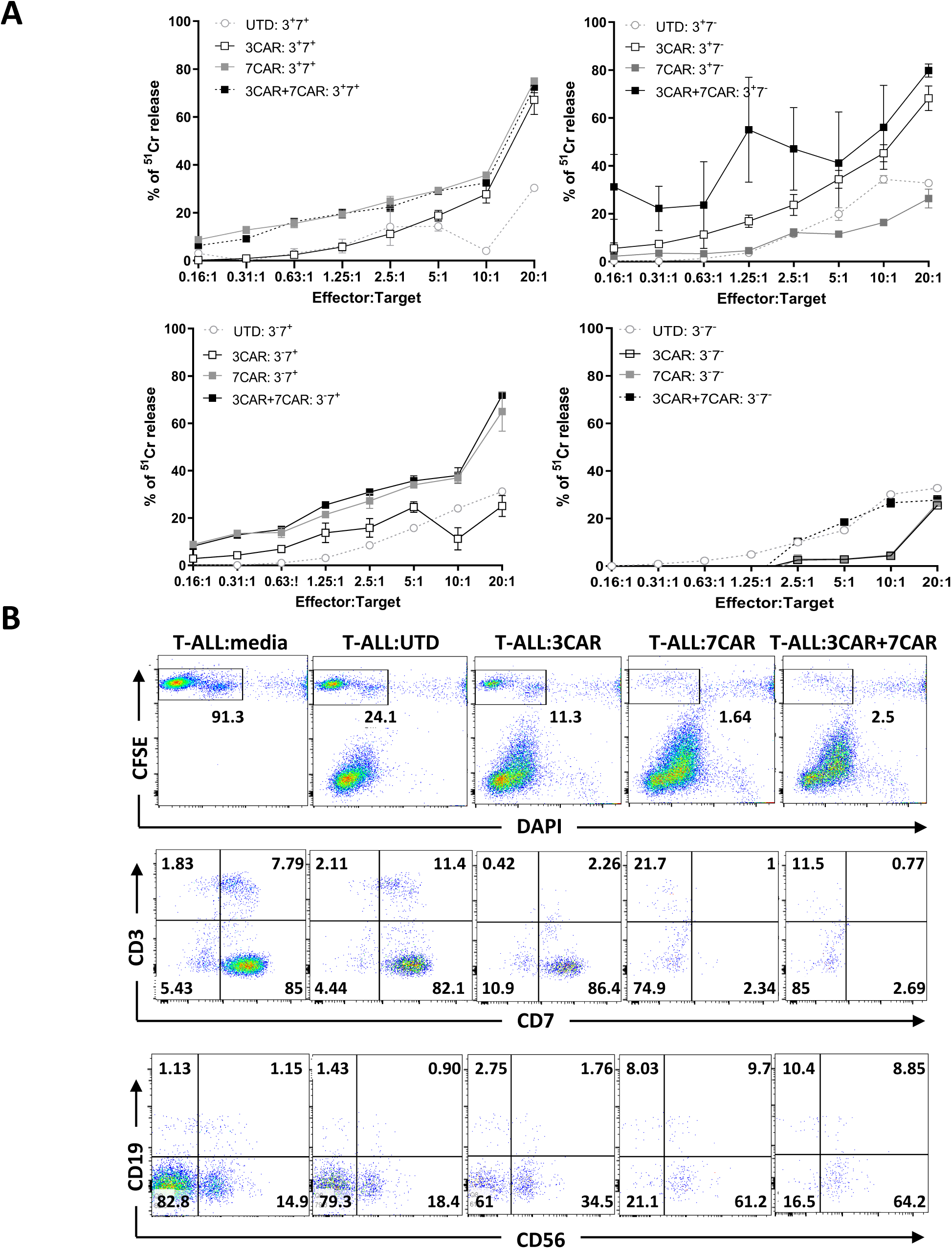
3CAR/7CAR T cells mediate potent killing of T-ALL cells *in vitro*. *In vitro* cytotoxicity of 3CAR and 7CAR cells against T-ALL cell lines and primary T-ALL targets. **A)** ^51^Cr labelled Jurkat T cells modified to express CD3^+^CD7^+^, CD3^+^CD7^-^, CD3^-^CD7^+^ or CD3^-^CD7^-^ were co-cultured with either 3CAR (white squares), 7CAR (grey squares), mixed 3CAR/7CAR (black squares) or untransduced (white circles) cells at an increasing ratio of effectors:targets (E:T). Error bars represent SEM of (n=3) technical replicates. **B)** Cytotoxic activity of 3CAR, 7CAR, mixed 3CAR/7CAR or untransduced primary T cell controls against primary patient T-ALL cells. Representative flow cytometry plots gated on CFSE^+^ live T-ALL tumour cells (top panel). Frequency of surface antigens CD3, CD7 (middle panel) and CD19, CD56, (lower panel) gated on CFSE+ live tumour cells.

Next the function of 3CAR and 7CAR base edited T cells was assessed against primary paediatric T cell acute lymphoblastic leukaemia (T-ALL). Flow based characterisation of n=7 primary T-ALL samples indicated high levels of cell surface CD7 in these patient samples, ranging from 83%-98%. In contrast, both surface and intracellular expression of CD3 were highly variable 1.5%-86% and 2%-92%, respectively, again highlighting the likely need to target multiple antigens in these subjects. A T-ALL sample where 9.6% of cells expressed surface CD3 and 92.8% expressed CD7, was loaded with fluorescent dye before culture with effector CAR T cells. Targets were almost depleted by 3CAR cells (residual 2.7% CD3) or 7CAR cells (residual 3.3% CD7) alone, and premixed 3CAR/7CAR T cells mediated almost complete elimination of target cells **(Figure 4B)**.

### Combinational anti-T cell CAR effects in humanised mice

*In vivo* anti-leukemic function of base edited CAR T cells was modelled in a humanised NOD/SCID/γc^-/-^ (NSG) xenograft model of leukaemic T cell inhibition. Mice (n=27) were engrafted with 1 x 10^7^ EGFP^+^LUC^+^ labelled Jurkat T cells modified to express CD3, or CD7, alone or in combination and 3 days later leukaemia establishment was confirmed by bioluminescent signalling **(Figure 5A & B)**. After 4 days, mice were inoculated with coBE3 edited 3CAR/7CAR T cells (after co-culture) or untransduced cells and leukaemic progression was monitored by serial bioluminescent imaging on days 7, 11, 14, 17, 21 and 24 **(Figure 5B & C)**. Within 7 days of 3CAR/7CAR effector infusion there was inhibition of luciferase signal in 3^+^7^+^ (n=5), 3^+^7^-^ (n=4) and 3^-^7^+^ (n=5) target groups, but not in the 3^-^7^-^ (n=5) cohort. By day 24, leukaemic burden in the 3^-^7^-^ infused group receiving 3CAR/7CAR effectors had increased >400-fold, comparable to leukaemic progression in the untransduced cohort (*ns P=0.36*) **(Figure 5C)**. This was further corroborated in the bone marrow of 3^-^7^-^ mice receiving mixed 3CAR/7CAR effectors through marked expansion of GFP^+^CD2^+^ tumour cells as was seen in mice receiving untransduced cells. In contrast, mice receiving premixed 3CAR/7CAR showed almost complete eradication of GFP^+^CD2^+^ T-ALL cells expressing one or both surface antigens CD3 and CD7 **(Figure 5D).**

**Figure 5.**
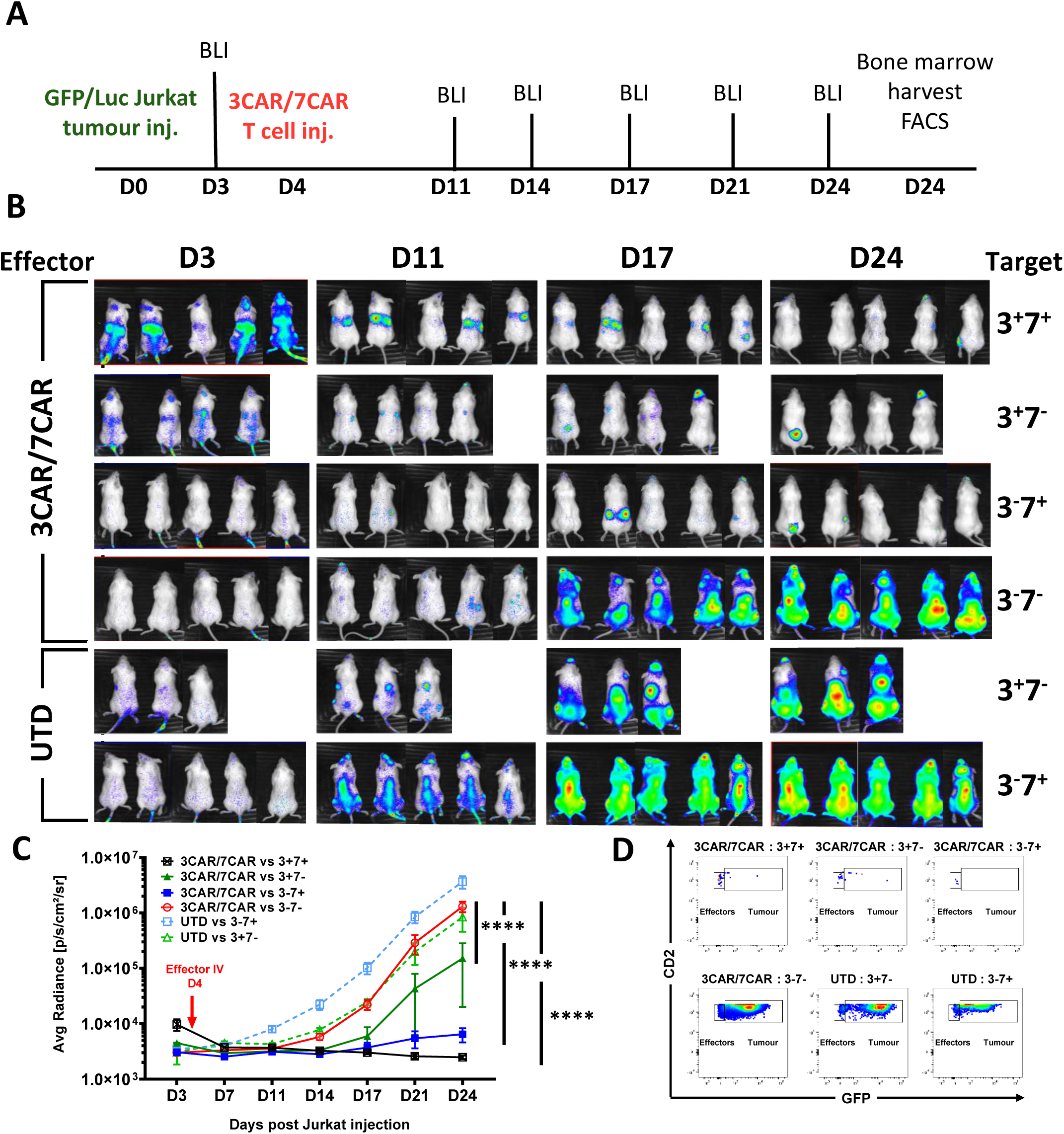
3CAR and 7CAR cells effectively clear T cell malignancy in vivo. **A)** Experimental timeline of GFP^+^LUC^+^ Jurkat tumour injection (Day 0) and effector T cell injection (Day 4) in n=27 NOD/SCID/γc^−/−^ (NSG) mice. Bioluminescent imaging (BLI) performed biweekly (Days 3-24). Organ harvest post mortem for flow-based characterisation (Day 24). **B)** NSG mice were infused with 1 x 10^7^ GFP+LUC+ Jurkat T cells modified to express mixed CD3 and/or CD7 surface antigens in groups of (n=5) CD3^-^CD7^-^, (n=4) CD3^+^CD7^-^, (n=5) CD3^-^CD7^+^ or (n=5) CD3^-^CD7^-^ and imaged on day 3 prior to infusion of 1 x 10^7^ TCR^-^CD7^-^ 3CAR/7CAR mixed effectors or untransduced (UTD) cells. Leukaemic progression monitored by serial BLI for 24 days and revealed disease progression in animals receiving untransduced T cells (3CAR^-^7CAR^-^) and in animals engrafted with antigen negative (CD3^-^CD7^-^) leukemia. **C)** Bioluminescence signal of each animal plotted as Average radiance [photons/sec/cm^2^/sr]. Each line represents a different experimental group and each point on the line the mean of each group. Error bars represent SEM. Area under the curve was calculated for each experimental group and values were compared using a one-way ANOVA with Tukey multiple comparison post-hoc*\*\*\*\*P<* 0.0001. **D)** Example of day 24 flow cytometry based detection in bone marrow of mCD11b^-^/hCD45^+^ effector T cells (CD2^+^GFP^-^) in in 3CAR/7CAR treated animals and residual leukemia (CD2^+^GFP^+^) in antigen negative and untransduced groups.

### Could promiscuous cytidine deamination corrupt CAR antigen-specificity?

Cytidine deamination by BE3 employs rat APOBEC1 fused to deactivated Cas9 nickase and a UGI moiety, and reports of guide independent activity both at the DNA and RNA level have raised concerns of promiscuous activity across the genome (21-23). Much of the published experience relates to experiments performed in cell lines following plasmid mediated expression of the editors and effects following mRNA mediated expression in primary cells may differ. Electroporation of BE mRNA into T cells was confirmed by Western blot to support transient protein expression (<48hrs) and should reduce the likelihood of off-target editing **(Supplementary Figure 3)**. Nonetheless, one area of concern is the possibility of deamination-mediated editing of the scFv antigen recognition elements that confer antigen specificity. Somatic hypermutation of antibody variable regions mediated by human cytidine deaminase in B cells has long been recognised as a mechanism to increase receptor diversity (24). Similar effects in CAR-T cells could inadvertently redirect cells away from desired targets leading to unknown specificity. To interrogate the integrity of CAR sequences following editing, total RNA was extracted from spCas9 and coBE3 mRNA modified T cells at 48 and 96 hours post transduction and again on day 14, at end of 3CAR and 7CAR production. cDNA was synthesised using universal primers flanking the CAR sequences and analysed by NGS for cytidine specific base conversion, focussing on the heavy and light antigen binding regions (ABRs) **(Figure 6)**. Quantification of cytidine transitions and transversions revealed that C->N occurrences were infrequent, shared by spCas9 and coBE3 edited samples, and generally ranged between 0%-2% within the ABRs **(Figure 6A)**. Only two changes were detected above this level, and only transiently, after 48hrs in 3CAR base edited T cells at positions P370 (4%) & P718 (7%) and had reduced at the end of production **(Figure 6C)**. Importantly there was minimal C->T transition across the entire CAR transgene in either spCas9 or coBE3 modified cells and the frequency of events detected 48hrs after mRNA delivery were broadly similar to those at the end of production on day 14 **(Figure 6B & C)**. Overall, there was no evidence to suggest sustained mutational corruption of antigen specificity in base edited CAR T cells.

**Figure 6.**
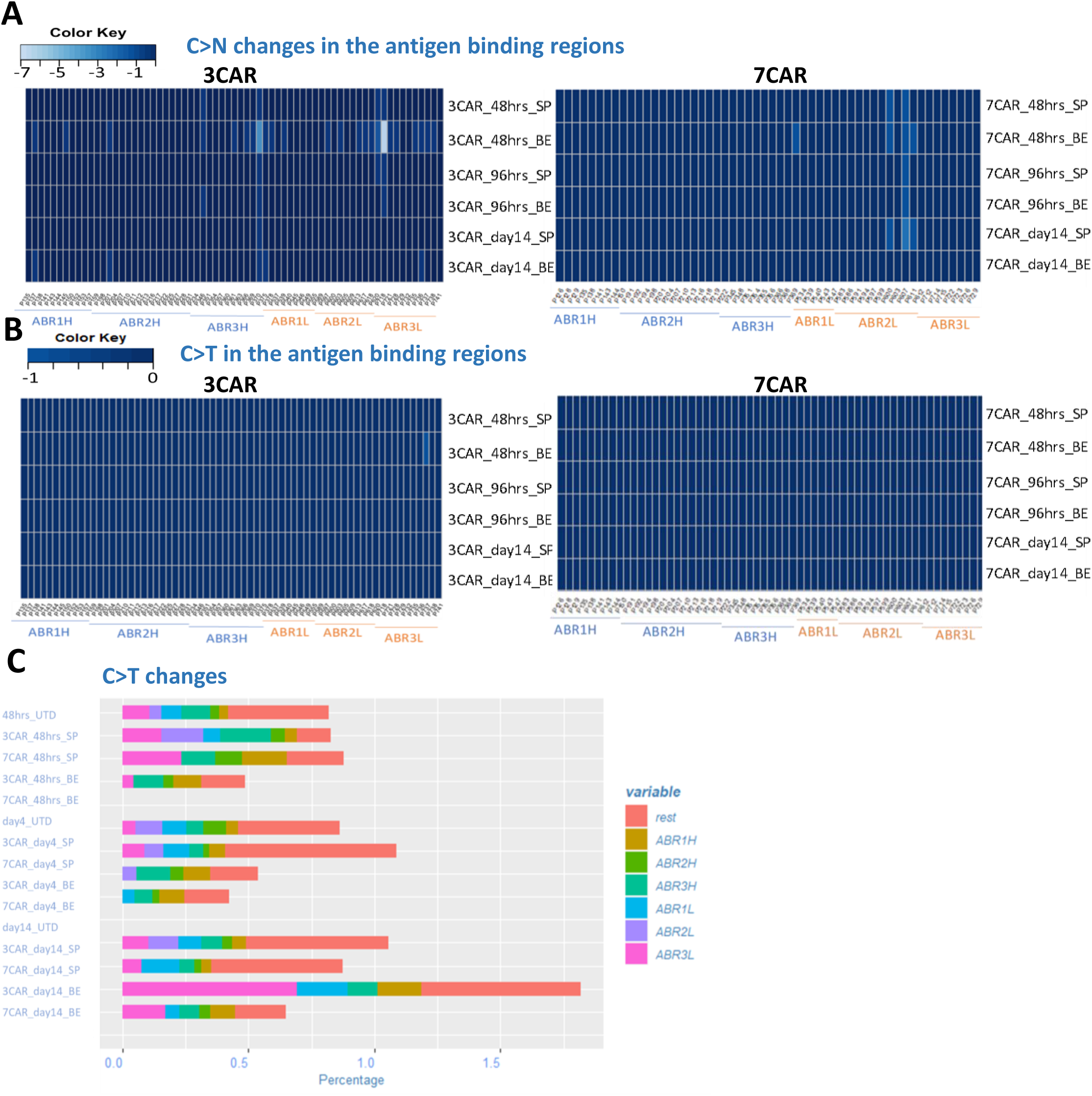
Cytidine deamination does not compromise integrity of antigen specificity of CAR sequences. Serial examination of 3CAR or 7CAR scFv RNA sequences 48hrs and 96hrs after electroporation with SpCas9 (SP3/SP7) or coBE3 (BE3/BE7) mRNA and again at end of production on d14. **A)** Amplicons of 3CAR (left) and 7CAR (right) vH and vL sequences with antigen binding regions (ABR) displayed mapped as a Heatmap in R using the *gplots* library for C>N conversion rates at the marked sites. **B)** 3CAR (left) and 7CAR (right) scFv ABR mapped as a Heatmap for C>T conversion rates. **C)** Stacked histogram showing <2% C>T changes in each region (ABR1H, ABR2H, ABR3H etc and outside the binding regions - (“rest”) in SpCas-CAR3, SpCas-CAR7, BE-CAR3, BE-CAR7 or untransduced (UTD) samples.

## Discussion

Engineered CAR T cell therapies for non-B cell malignancies are proving challenging, with particular difficulties arising when target antigens on target populations are also present on T cells. We have previously described strategies to address fratricide for the generation of CAR T cells against CD3, a definitive T cell marker present on certain T cell lymphomas and a proportion of T-ALL (9). Electroporation of TALEN mRNA was used to prevent assembly and expression of the multimeric TCRαβ/CD3 complex. The resulting T cells were not only insensitive to the 3CAR but were simultaneously rendered TCRαβ negative, and thus non-alloreactive and incapable of mediating graft versus host disease if derived from an unmatched healthy donor. Approaches using ‘off-the-shelf’ universal CAR T cells against CD19 are already under investigation, including cells with additional genomic edits to address the risk of host mediated rejection (25, 26). CAR19 T cells edited at the TRAC and CD52 locus using TALEN technology are undergoing multicentre testing, and similar approaches using homing endonuclease editing or CRISPR/Cas9 are under investigation (17, 18, 27). An ability to pre-manufacture universal CAR T cells opens up the possibility of generating cell banks with different specificities, which could then be used in combination as a strategy to address incomplete or internalised target expression and antigen escape. In the context of CAR T cells targeting T cell malignancies, a time-limited application is envisaged to allow recovery of a functional T cell compartment, derived either from autologous, antigen-negative precursors or donor derived stem cells following allo-SCT. Suitable additional antigens for targeting T cell malignancies include TCRαβ, CD2, CD5 and CD7. A strategy to target TCRαβ on T cell lymphomas is in trial and relies on discriminatory targeting of TRBC1 or TRBC2, but would not be suitable to combine with CD3ε given their linked multimeric expression (28). CD2 is expressed on mature T cells and has a role in adhesion and co-stimulation upon interaction with its ligand CD58 (LFA-3) (29). Alefacept, a recombinant LFA-3/IgG1 fusion protein was previously investigated as an immunosuppressive anti-CD2 treatment for psoriasis, and was noted to selectively deplete memory T cell subsets (30, 31). In depth functional studies of CD2 in animals is sparse, mainly because LFA-3 in mice is absent and normal lymphocyte compartments with CD2 knockouts may not be representative (32). Monoclonal antibodies against CD5 have also been investigated and shown encouraging results against cutaneous T-cell lymphoma (CTCL) and T-ALL (33). The molecule is downregulated in activated T cells and anti-CD5 CAR T cells co-expressing a CD28 costimulatory domain exhibited only transient fratricide (34), but 4-1BB coding constructs exhibited impaired expansion of CD5 CAR T cells and higher levels of fratricide were attributed to tumour necrosis factor (TNF) receptor associated factor (TRAF)-mediated upregulation of intracellular adhesion molecule 1 (ICAM1) (35). We have focussed on CD7, which has previously been investigated as a target for monoclonal antibody immunotoxin therapy against T cell malignancies (36). Recently the introduction of strategies to inhibit expression on T cells has enabled the anti-CD7 scFv binding domains to be employed in CAR approaches. Gomes-Silva et al reported combining CRISPR/Cas9 editing to eliminate CD7 in T cells expressing anti-CD7 CARs and have modelled strategies in humanised mice against T cell lymphoma and acute myeloid leukaemia (15). In addition, an alternative approach using ‘protein expression blockade’ has employed an anti-CD7 scFv moiety coupled to endoplasmic reticulum/Golgi retention elements to anchor the protein and prevent surface expression (13). Of note, these recent reports have found no evidence of impaired effector cytotoxicity after disruption of CD7, and CD7 deficient mice appear to retain normal lymphoid compartments (37). The cytoplasmic tail of CD7 activates phosphoinositol signalling of CD7 in T cells and may play a role in co-stimulation (38). Secreted epithelial protein K12/SECTM1 is the ligand for CD7 and is an interferon inducible protein encoded at the continuous genetic locus for CD7, indicating feedback through transcriptional regulation may be in operation (39, 40). Given that CAR T cells disrupted for CD7 retain potent functional activity, it appears that CD7 signalling, and by proxy SECTM1 activity, may be redundant for anti-tumour function.

In consideration of allogeneic cells, Cooper et al described a strategy to generate ‘universal’ donor T cells with simultaneous TCR/CD3 and CD7 disruption using CRISPR/Cas9 editing to generate cells expressing their anti-CD7 CAR (14). The cells were investigated against T cell lines and primary T cell malignancies, and included studies in chimeric xenografted mice. The issue of possible translocations from multiplexed editing was not investigated, but we report a similar editing frequency and found readily detectable and quantifiable events between sites of guide activity at the *TRBC* and *CD7* loci. The frequency was consistent with past experiences with TALENs and CRISPR/Cas9. In contrast, upon switching to BE3, and using the same sgRNA guides, these events were virtually eliminated.

Given that additional genome edits would be required to confer resistance to Alemtuzumab or disrupt expression of checkpoint pathways such as PD1, the frequency of additional translocation events would become problematic. We have found that additive co-delivery of sgRNAs against CD52 conferred high levels of triple knockout (TCRαβ/CD7/CD52) with no alternation in phenotype or cytotoxic responses (data not shown) and anticipate transformation risks will be mitigated compared to conventional Cas9. However, definitive experiments to prove an enhanced safety profile in T cells are difficult to conceive given that mature T cells are not readily transformed. Similarly, it is not readily possible to quantify adverse impacts that may arise from ectopic deamination activity, either as a result of guide directed BE effects at possible off target sites, or as a result of promiscuous, guide-independent, APOBEC activity on DNA or RNA. One potentially serious consequence of aberrant base conversion would be changes to antigen receptor sequences, in particular if involving regions defining antigen binding specificity. Interrogation of RNA collected at serial timepoints over a 14 day period from electroporation of BE mRNA until end of production found no evidence of deamination mediated transition, transversion (or other aberrations) within the antigen binding frameworks of the scFv for 3CAR or 7CAR. This finding is reassuring, but in any case unwanted activity has been greatly reduced in recent BE variants.

The processes described here are readily scalable and similar techniques have already been deployed in a compliant manner for CRISPR/Cas9 edited CAR19 T cells. The issue of stringently depleting residual TCRαβ T cells remains critical for non-HLA matched donor T cells, and the Prodigy device supports automated microbead depletion with <1% TCRαβ expression in the final products. This step is retained for anti-T cell CARs even though CAR mediated ‘self-enrichment’ arises during production and will ensure strict specification limits. It is anticipated that sufficient production for both 3CAR and 7CAR will be achievable from a single non-mobilised peripheral blood leukapheresis to support dozens of therapeutic doses, offering a route to affordable and accessible cell therapy.

## Materials and Methods

### Cell lines

Jurkat cells (acute T cell leukemic cell line from ATCC) were modified to express CD3^+^CD7^+^ CD3^+^CD7^-^, CD3^-^CD7^+^ and CD3^-^CD7^−^ by spCas9 mRNA-based disruption of TCR and/or CD7 using TRBC sgRNA and CD7 sgRNA and maintained in culture in RPMI-1640 (Thermo Fisher Scientific) supplemented with 10% FBS (Millipore Sigma) **(Supplementary Figure 4)**. For *in vivo* experiments cells were stably transduced with a 3^rd^ generation pCCL-PGK-EGFP-LUC lentiviral vector and were sorted on a MoFlo XDP (BD) for GFP expression prior to banking.

### SpCas9 and coBE3 genome editing

Synthetic sgRNAs were manufactured by Synthego (California, US) by automated solid-phase synthesis with 2’-O-methyl 3’ phosphorothioate modifications. Single guide RNA containing a 20 nucleotide protospacer with an 80 nucleotide CRISPR scaffold were eluted in nuclease-free Tris-EDTA buffer

sgRNA guide sequences for TRBC 1 and 2 and CD7 were selected for compatibility with both SpCas9 and coBE3.

CleanCap^®^ Cas9 mRNA encoded SpCas9 was supplied by Trilink US. Custom made codon optimised BE3 (coBE3) was supplied as CleanCap (Trilink US) with a Cap 1 structure, and polyadenylated to increase expression and stability.

### Generation of 3CAR and 7CAR vectors

3CAR expressing OKT3 monoclonal antibody scFv was previously described (9). 7CAR was derived by codon optimization (GeneArt) of variable heavy chain and variable light-chain antigen binding elements of the anti–human CD7 murine hybridoma, 3A1e sequence (15).The scFv was fused to a CD8 transmembrane domain and to activation domains derived from 41BB and CD3ζ. The resultant 7CAR construct was cloned into a lentiviral vector (pCCL) backbone under the control of a PGK promoter. Vector stocks pseudotyped with a vesicular stomatitis virus glycoprotein (VSV-G) envelope were generated in 293T cells (ATCC) and yielded titres >5 × 10^8^ transducing units/ml after ultracentrifugation (41).

### Generation of fratricide resistant 3CAR and 7CAR effectors

PBMC were obtained from healthy donors and were cultured in 48-well plates at a density of 1 × 10^6^/ml in TexMACS (Miltenyi Biotec), 3% human serum (Seralab) +20 ng/ml human recombinant IL-2 (Miltenyi Biotec) and activated with TransAct reagent (Miltenyi Biotec). Activated T cells were electroporated with TRBC sgRNA (Synthego), CD7 sgRNA (Synthego) and codon optimised BE3 mRNA (TriLink Biotechnologies) in a Lonza 4D or LV nucleofector (Lonza, Slough, UK) and cultured at 30°C 5% CO_2_ overnight before returning to 37°C. Cells were transduced with 3CAR or 7CAR lentiviral vector preparations at an MOI of 5 the following day. Cells were cultured in G-Rex 10 or G—Rex 100 chambers as per manufacturer’s instructions (Wilson Wolf, MN, USA).

### Flow cytometry

Cells were stained with the following primary anti-human antibodies: mouse anti-human CD2 (clone LT2), mouse anti–human CD3 (clone BW264/56), mouse anti–human TCRαβ (clone BW242/412), mouse anti–human CD4 (clone VIT4), mouse anti–human CD8 (clone BW135/80), mouse anti–human CD19 (clone LT19), mouse anti–human CD56 (clone AF12-7H3), (all from Miltenyi Biotec), mouse anti-human CD7 (clone M-T701) (BD), mouse anti-human CD7 (clone 3A1e) (MyBioSource), mouse anti-human CD7 (clone MEM186) (Abcam), mouse anti-human CD7 (clone 6B7) (BioLegend). To assess the efficiency of 3CAR and 7CAR transduction, cells were stained using a Biotin-SP (long spacer) AffiniPure Fab Fragment Goat Anti-Mouse IgG fragment-specific antibody (catalogue 115-066-072; Jackson Immunoresearch, Stratech Scientific Limited) followed by Streptavidin-APC (catalogue 405207, BioLegend) or Streptavidin-FITC (catalogue 405202, BioLegend) or CD7 Protein, Human, Recombinant (His Tag) (catalogue 11028-H08H, Sino Biological) followed by Anti-6X His tag^®^ antibody [AD1.1.10] (DyLight^®^ 650) (catalogue ab117504, Abcam). Cells were acquired on a Cyan or on a BD LSRII (BD Biosciences), and analysis was performed using FlowJo v10 (TreeStar Inc.).

### Molecular quantification of on-target genome editing

Genomic DNA extraction was performed using DNeasy Blood and Tissue Kit (69504, QIAGEN) and a PCR reaction designed to amplify 400-800bp over the protospacer binding site for *TRBC* and *CD7* genomic loci. PCR products were gel extracted and sent for Sanger sequencing (Eurofins Genomics). Resulting Sanger sequencing data was analysed using TIDE (https://tide.nki.nl/), to measure the frequency of indels, at the predicted SpCas9 scission site **(Supplementary Figure 2B)**. When analysing C>T conversion rates produced by coBE3, EDITR software was used (https://moriaritylab.shinyapps.io/editr_v10/).

### Translocation detection and quantification

To detect potential chromosomal translocations following multiplexed editing, primers flanking genome editing region at *TRBC* locus within chromosome 7 q-arm and CD7 locus within chromosome 17 q-arm were designed amplifying translocations T1-T4. *TRBC* Fwd - *CD7* Fwd (T1), *TRBC* Rev – *CD7* Fwd (T2), TRBC Rev – CD7 Rev (T3) and TRBC Fwd – CD7 Rev (T4) primer combinations were used to PCR amplify SpCas9 or coBE3 edited mixed 3CAR/7CAR or untransduced DNA products. Plasmids carrying the predicted fusions were synthesised as positive controls (GeneArt).

A ddPCR (BioRad) was used for the accurate quantification of the TRBC:CD7 translocations. Each sample was run as a duplexed assay consisting of translocation-specific primer pairs (designed for each translocation for a maximum of 230bp product) and probes as well as an internal (albumin) reference primer and probe set. All primers and probes were ordered from IDT. PCR reactions were set up using the ddPCR Supermix for Probes, no dUTP (BioRad). Droplets were generated and analysed using the automated QX200 AutoDG Droplet Digital PCR System by BioRad. Analysis was done on Quantasoft (BioRad) and in R using an optimised version of twoddPCR (https://github.com/CRUKMI-ComputationalBiology/twoddpcr/) for the purpose of this study.

### Detection of Cas9 protein

Cell pellets (1 x 10^6^ cells) were acquired at 12, 24, 48, 72 and at day 7 post electroporation with coBE3 mRNA. Total protein was quantified by a bicinchoninic acid (BCA) assay. Western blot was run using 25μg total protein per sample. 16.5ng Alt-R^®^ S.p. HiFi Cas9 Nuclease V3 (1081061, IDT, Leuven, Belgium) was used as positive control. Membrane was blotted with mouse anti-CRISPRCas9 antibody (ab191468, abcam, Cambridge, UK) at a 1:1000 dilution in 3% milk overnight at 4°C before incubation with secondary HRP-linked sheep anti mouse (NA931-1ML, GE Healthcare Life Sciences, Buckinghamshire, UK) at a 1:3000 dilution in 5% milk for 1 hour at room temperature. Protein was visualized by chemiluminescence using Pierce ECL western blotting substrate (32106, ThermoFisher Scientific).

### Chromium release assay of *in vitro* cytotoxicity

Cytotoxic activity of 3CAR and 7CAR cells was assessed by ^51^Cr release assay. For this, 5 × 10^3 51^Cr-labeled CD3^+^CD7^+^, CD3^+^CD7^-^, CD3^-^CD7^+^ and CD3^-^CD7^-^ Jurkat cells, were incubated with either 3CAR, 7CAR, mixed 3CAR/7CAR or untransduced control effector cells at increasing effector:target ratios (E:T ratios) in 96-well microplates for 4 hours at 37°C. Supernatant was subsequently harvested and mixed with OptiPhase HiSafe 3 (PerkinElmer) scintillation fluid and incubated at RT for 16hrs. ^51^Cr release was then measured in a microplate scintillation counter (Wallac 1450 MicroBeta TriLux). Specific lysis was calculated using the formula [(experimental release - spontaneous release)/(maximum release - spontaneous release) x 100].

### *In vivo* antitumor activity

Ten-week-old female NOD/SCID/γc^−/−^ NSG mice (Charles River, strain: NSG [005557] from The Jackson Laboratory), were inoculated i.v. with 1 × 10^7^ CD3^+^CD7^+^, CD3^+^CD7^-^, CD3^-^CD7^+^ and CD3^-^CD7^−^Jurkat T cell tumour targets by tail vein injection on day 0. The Jurkat T cell targets had been stably transduced to express EGFP^+^LUC^+^ and, following CRISPR/Cas9-mediated TRBC1/2 and/or CD7 disruption, had been sorted for CD3^+^ and CD3^−^, CD7^+^ or CD7^-^ expressers. Tumour engraftment was confirmed by *in vivo* imaging of bioluminescence using an IVIS Lumina III *In vivo* Imaging System (PerkinElmer, live image version 4.5.18147) on day 3 and mice were further injected on day 4 with either 1 × 10^7^ untransduced T cells, or 1 × 10^7^ mixed 3CAR/7CAR T cells. Analysis of tumour clearance was performed by serial bioluminescent imaging on days 3, 7, 11, 14, 17, 21 and 24, and processing of BM for the monitoring of tumour progression vs. clearance was carried out on day 24. BM samples were processed by a RBC lysis, followed by staining for flow cytometry. All animal studies were approved by the UCL Biological Services Ethical Review Committee and licensed under the Animals (Scientific Procedures) Act 1986 (Home Office, London, United Kingdom).

### Primary T-ALL blasts

Primary T-ALL patient samples were provided by the Bloodwise Childhood Leukaemia Cell Bank, Stockport, United Kingdom. To phenotype, patient cells were stained with the following primary antibodies: mouse anti–human CD3 (clone BW264/56; Miltenyi Biotec), mouse anti–human CD7 (clone M-T701) (BD Biosciences).

For the FACS-based killing assay, 10^5^ T-ALL patient target cells loaded with CFSE dye (CellTrace, Invitrogen) were incubated for 24 hours at 1:1 ratio with either 3CAR, 7CAR, mixed 3CAR/7CAR effector T cells or control untransduced T cells.

Study approval. Human sample collections were approved by University College London (UCL) ethics committee and collected with written consent. Leukaemia samples were provided by the Childhood Leukaemia Cell Bank.

### 3CAR and 7CAR targeted RNA sequencing

Samples were treated with either SpCas9 or coBE3 then subsequently transduced with 3CAR or 7CAR. RNA was extracted from samples at different timepoints using the RNeasy mini kit (Qiagen). DNase treatment was performed with RQ1 DNase (Promega) followed by reverse transcription using a primer specific to both the 3 and 7 CAR. The resulting cDNA was then purified using the Qiagen PCR purification kit and amplified using primers specific to 3 and 7 CAR sequences. PCR products were size selected using Promega beads. Once quantified by Qubit fluorometry and checked using Tapestation electrophoresis, libraries were prepared for sequencing following the Illumina Nextera XT protocols. Barcoded libraries were pooled, denatured and finally, sequenced on a MiSeq with a 500-V2 cartridge. Fastq files were downloaded using basespace Illumina and analysed with a homebrewed pipeline including steps for quality check (fastp), trimming (TrimGalore), removing PCR duplicates (Mark Duplicate reads), alignment to reference(s) (Bowtie2) but also investigation for minor allele frequencies (Naïve Variant Caller) and indels (Pindel). Sequences were visualized on Integrative Genomics Viewer (IGV). All figures were produced in Rstudio.

### Statistics

For comparison of in vivo study groups, area under the curve (AUC) was calculated for the grouped data. Values were compared using a one-way ANOVA with Tukey multiple comparison post-hoc. Values are presented as mean percentages of 3 or more samples with SEM or SD. In all experiments, P < 0.05 was considered statistically significant. All statistical analysis was performed using GraphPad Prism software version 8.0.

## Disclosures

WQ holds interests unrelated to this project in Autolus Ltd.

WQ received unrelated research funding from Cellectis, Servier, Miltenyi, Bellicum.

## Funding

Supported by the Wellcome Trust (215619/Z/19/Z), NIHR (RP-2014-05-007), NIHR Blood and Transplant Research Units (BTRU) and Great Ormond Street Biomedical Research Centre (IS-BRC-1215-20012), Bloodwise and Children with Cancer (2014/171).

The views expressed are those of the author(s) and not necessarily those of the NHS, the NIHR, or the Department of Health.

## Figure Legends

**Supplementary Figure 1. 7CAR expression results in partial epitope masking of CD7 surface antigen.** Flow cytometry plots of untransduced (UTD) or 7CAR transduced PBMCs without CD7 disruption stained for CAR transduction and CD7 expression using four distinct anti-CD7 antibody clones.

**Supplementary Figure 2. SpCas9 3CAR and 7CAR primary T cells evade fratricide during production**.

**A)** Representative phenotypic analysis of CD3 and CD7 surface antigen expression (top panel) and CAR transduction (bottom panel) of CD3/CD28 activated PBMC cells elcetroporated with TRBC and CD7-targeting sgRNA and spCas9 mRNA and subsequently transduced with 3CAR or 7CAR lentiviral vectors at MOI 5. Self-enrichment effects following co-culture of 3CAR and 7CAR products results in enriched TCR^-^CD7^-^ 3CAR/7CAR cells (red box). **B)** Representative TIDE output of Sanger sequencing results from mixed 3CAR/7CAR DNA PCR amplified using TRBC or CD7 forward and reverse primers. Percentage of target indels shown by red bars. **C)** In vitro cytotoxic activity of spCas9 or coBE3 edited 3CAR, 7CAR cells or untransduced negative control effector T cells against ^51^Cr loaded CD3^+^CD7^+^ Jurkat T cell targets at an increasing effector:target (E:T) ratio. Points represent mean of n=3 experimental triplicates and error bars represent SEM.

**Supplementary Figure 3. Base editor protein dilution following primary T cell culture.**

Cell lysates from primary T cells transduced with a Terminal-TRAC-CAR19 lentiviral vector at MOI 5 and exposed to coBE3 mRNA by electroporation were collected 12, 24, 48, 72 hrs post culture and again on day 7 of culture. Untransduced cell lysate was also collected at day 7 and used as a negative control. 20μg of BCA-normalised total protein was loaded on a 4%-15% SDS-PAGE alongside positive spCas9 protein control (15.6ng). Membrane was probed with anti-CRISPR-cas9 primary and sheep anti-mouse-HRP secondary antibody. Cas9 positive bands detected at 205kDa. Membrane was probed with anti b-actin primary antibody and sheep anti-mouse-HRP secondary antibody as a marker of loading detected at 40kDa.

**Supplementary Figure 4. Jurkat T cells modified for to express CD3 and/or CD7 surface antigen.**

Jurkat T cells electroporated with spCas9 mRNA and TRBC and/or CD7 targeting sgRNAs and sorted for CD3^+^CD7^+^, CD3^+^CD7^-^, CD3^-^CD7^+^ and CD3^-^CD7^-^. Flow cytometry of CD3 and CD7 expression of edited and sorted Jurkat T cells confirming surface antigen expression.

