## Supplementary figures and images for "Base-edited CAR T Cells for combinational therapy against T cell malignancies"

### Supplementary Figures S1, S2, S3, S4

#### Supplementary Figure 1

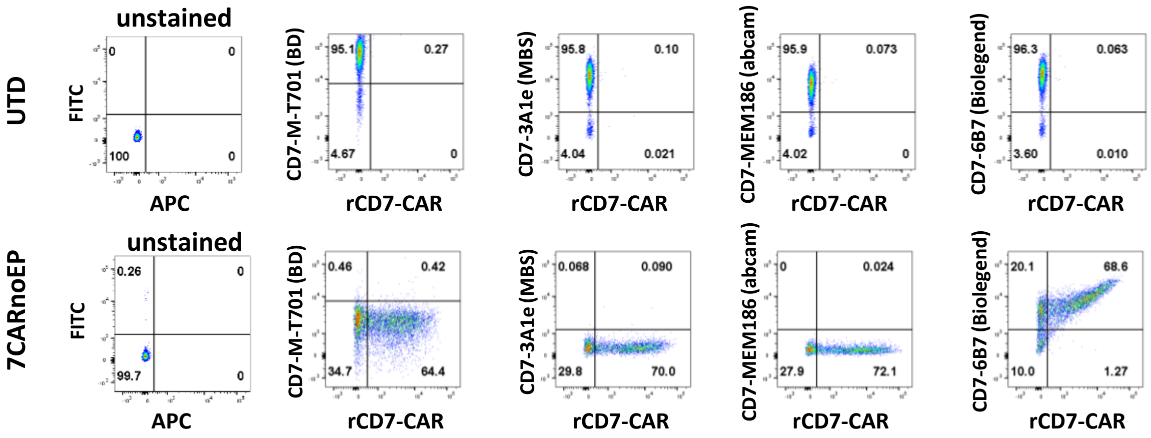

#### Supplementary Figure 2

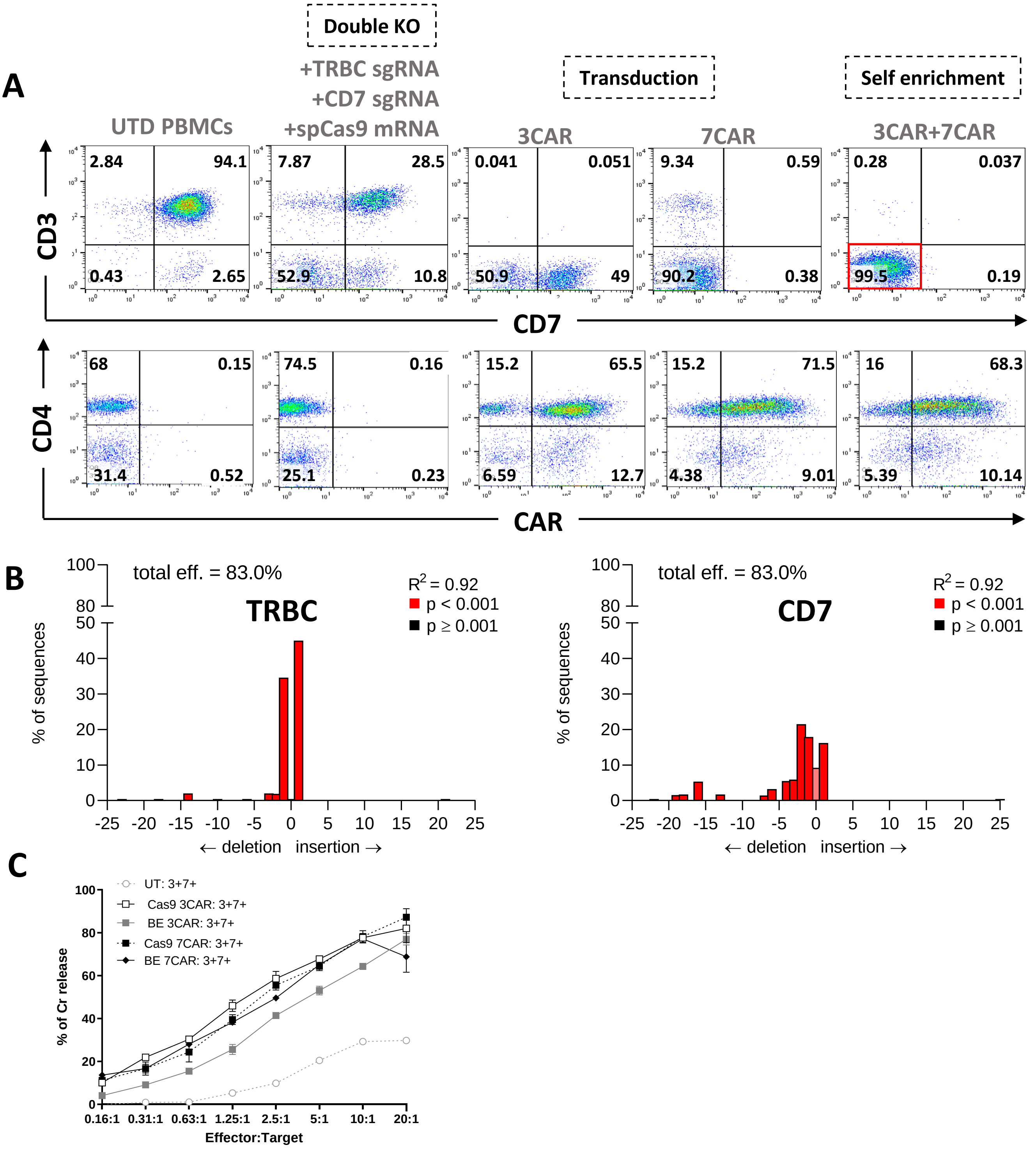

#### Supplementary Figure 3

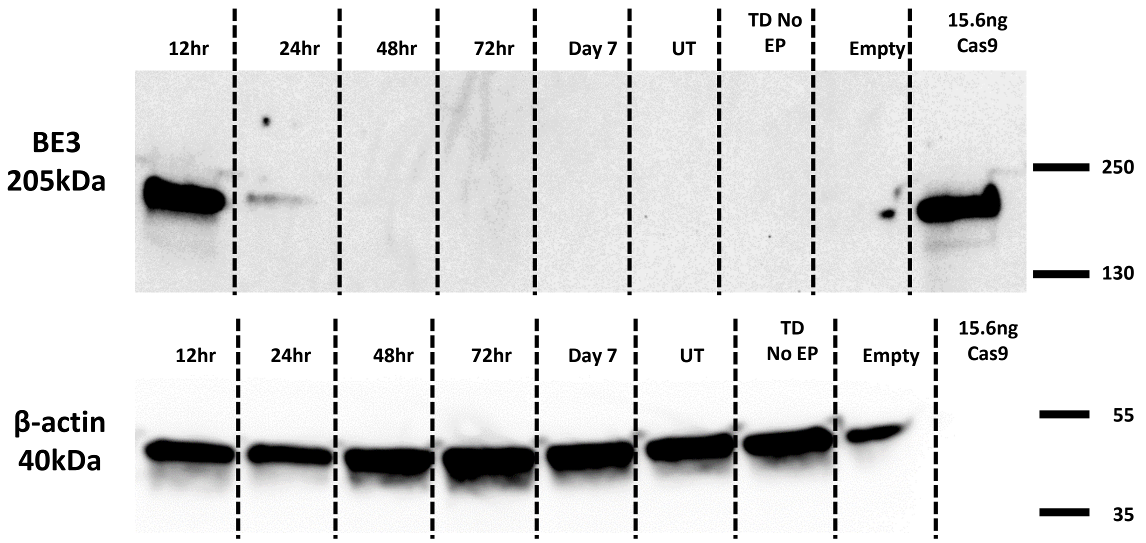

#### Supplementary Figure 4

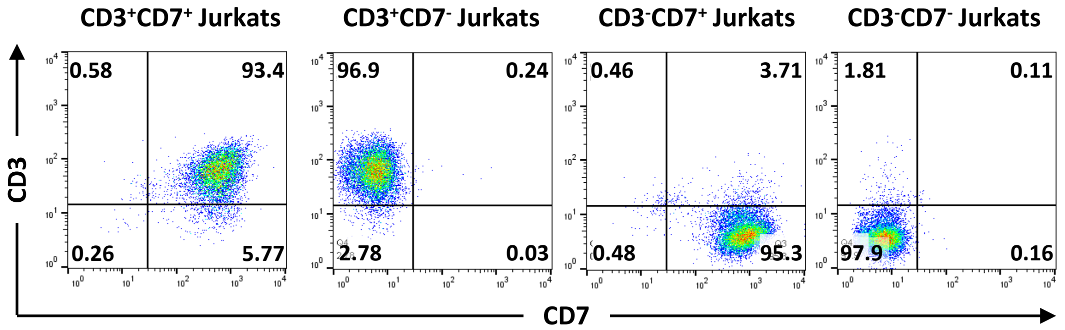
